## Supplementary figures and images for "Hidden Markov Models Reveal Behavioral State Dynamics in Depth-Related Locomotion in Mice"

### Supplemental Figure 1

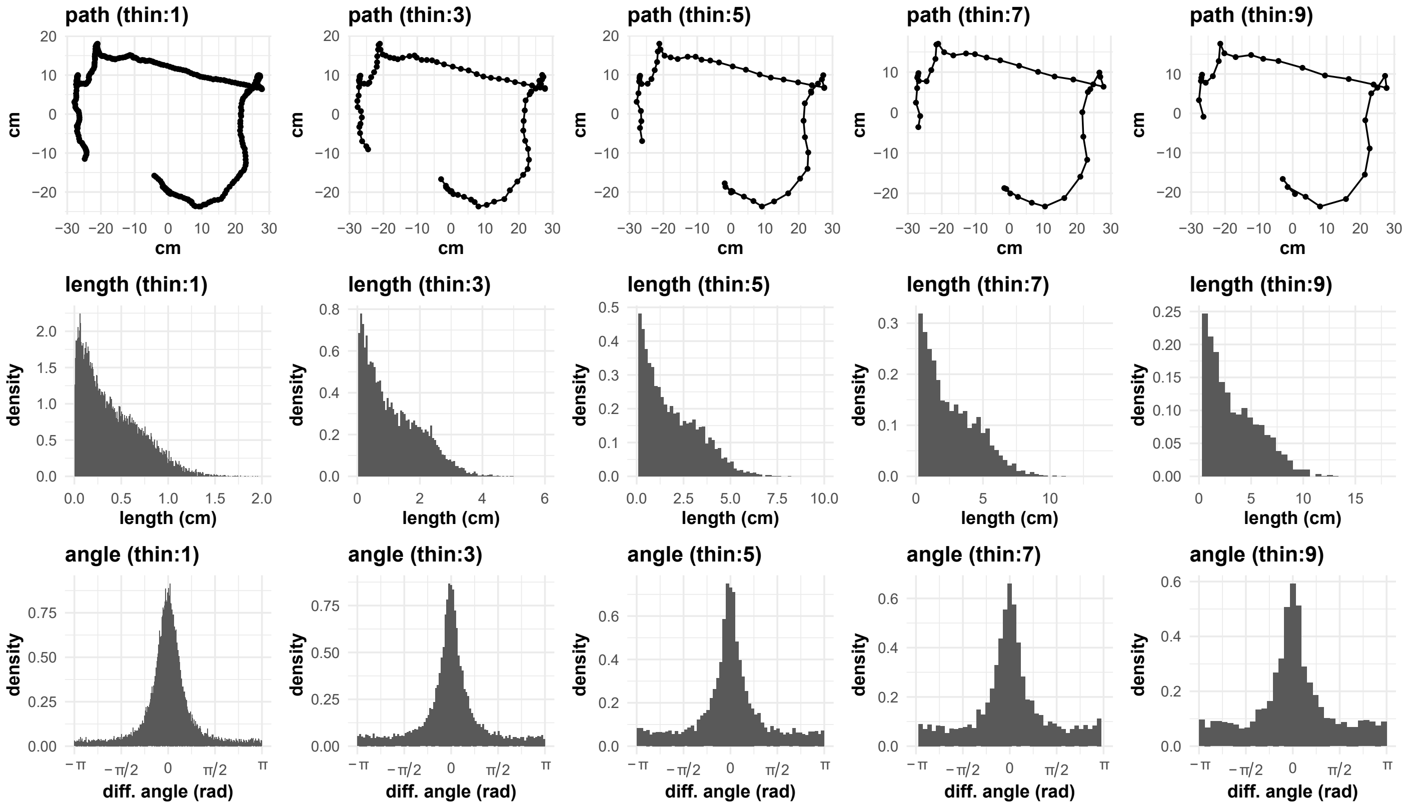


#### Supplemental Figure 1

### Supplemental Figure 2

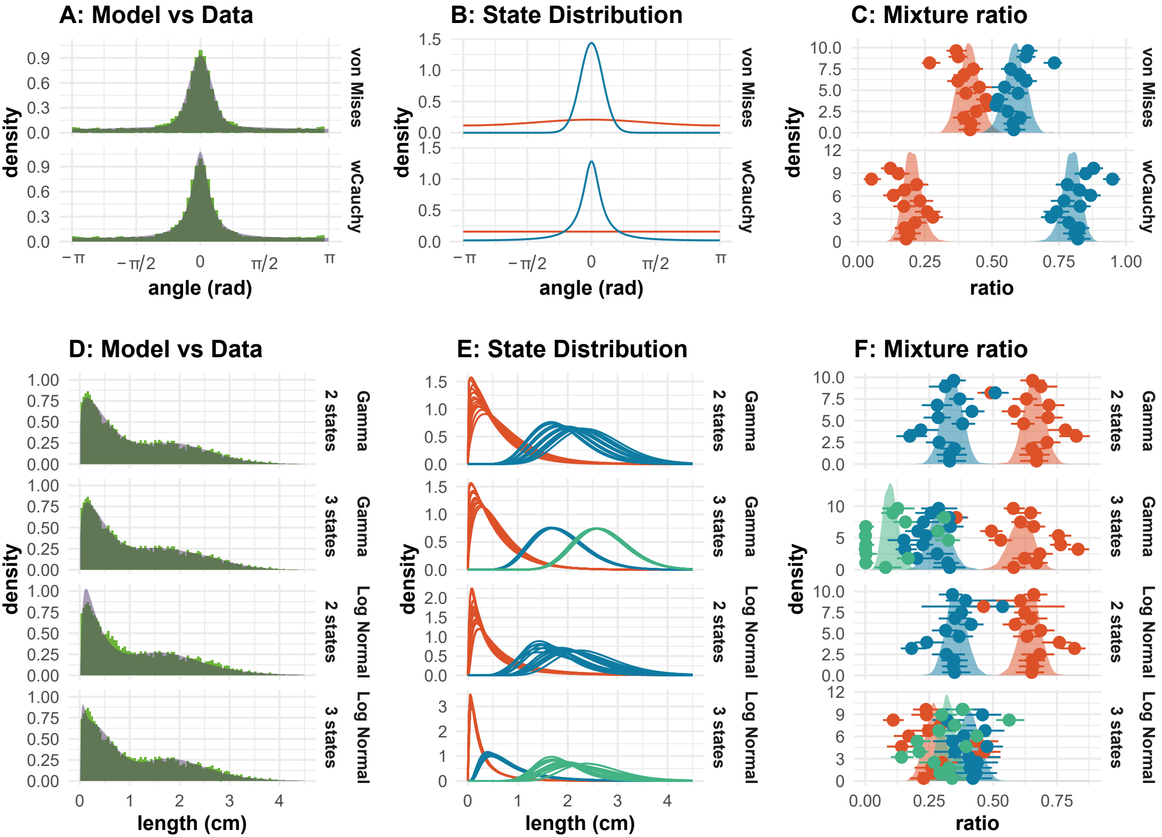


#### Supplemental Figure 2

### Supplemental Figure 3

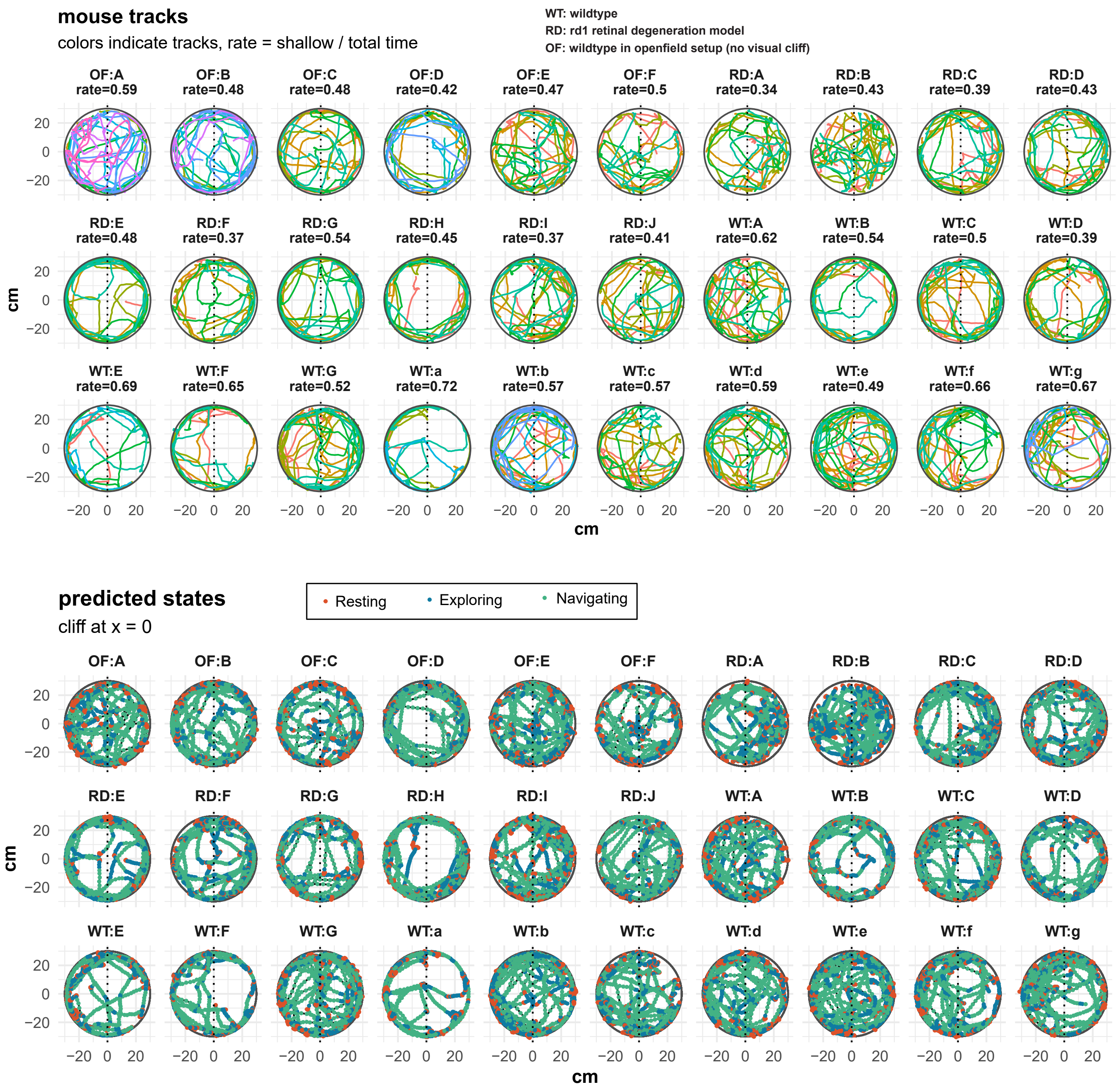


#### Supplemental Figure 3

### Supplemental Figure 4

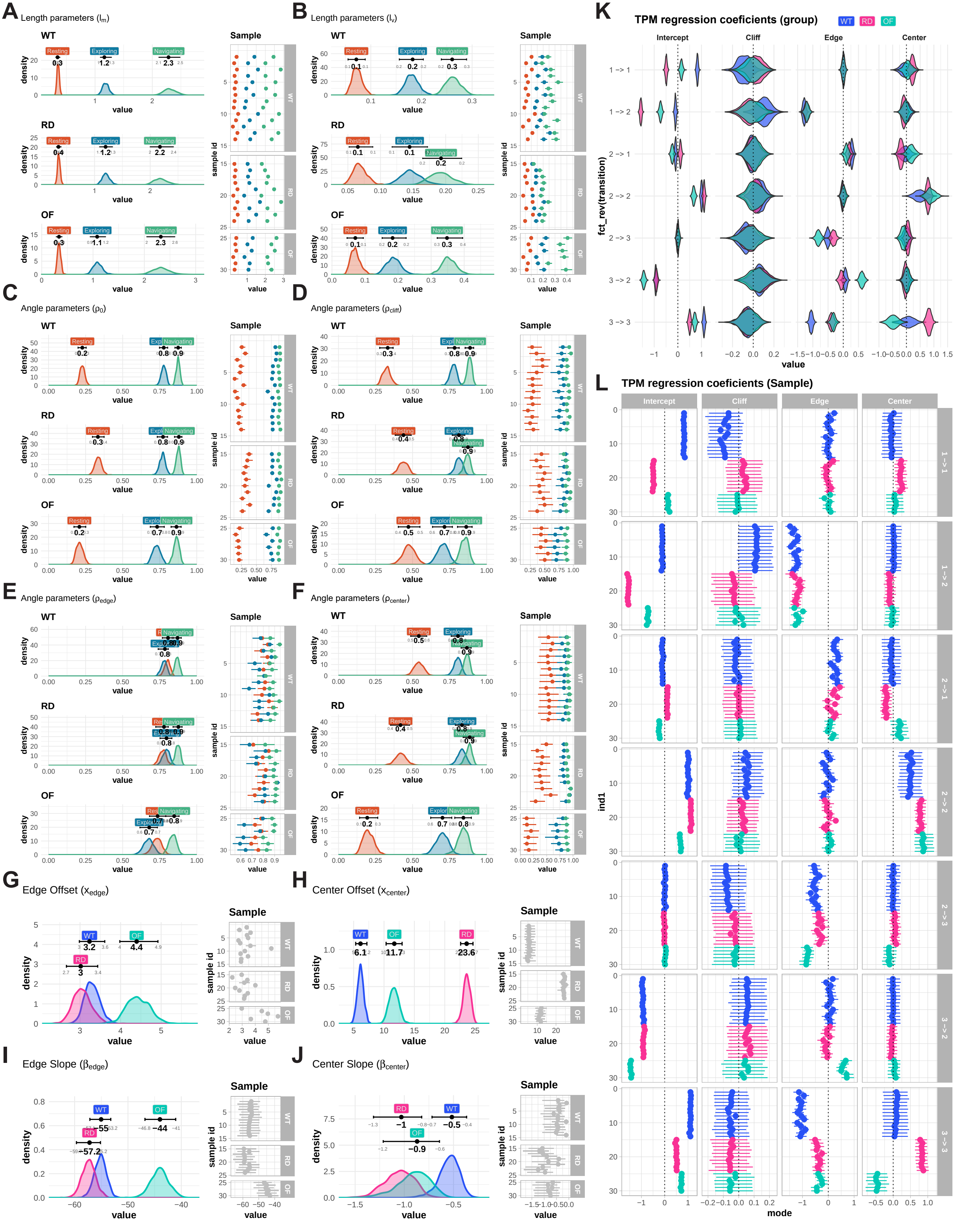


#### Supplemental Figure 4

### Supplemental Figure 5

##### Size

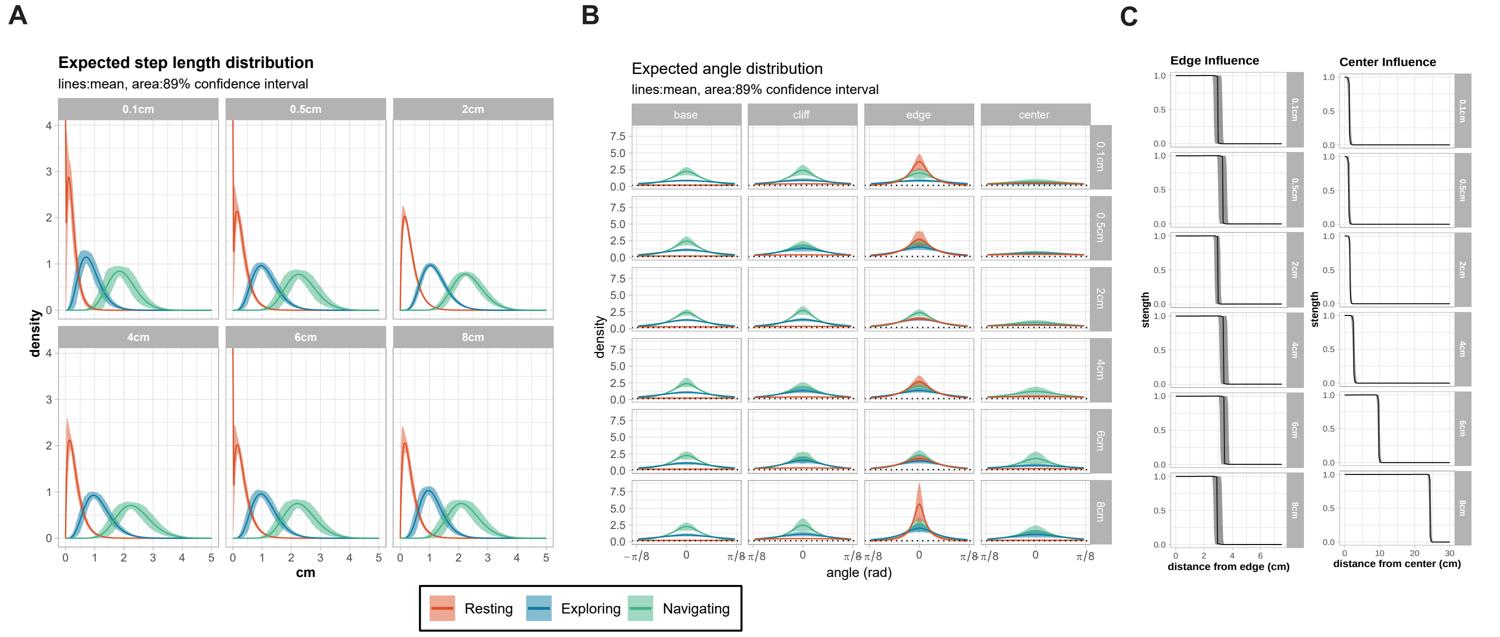


##### Contrast

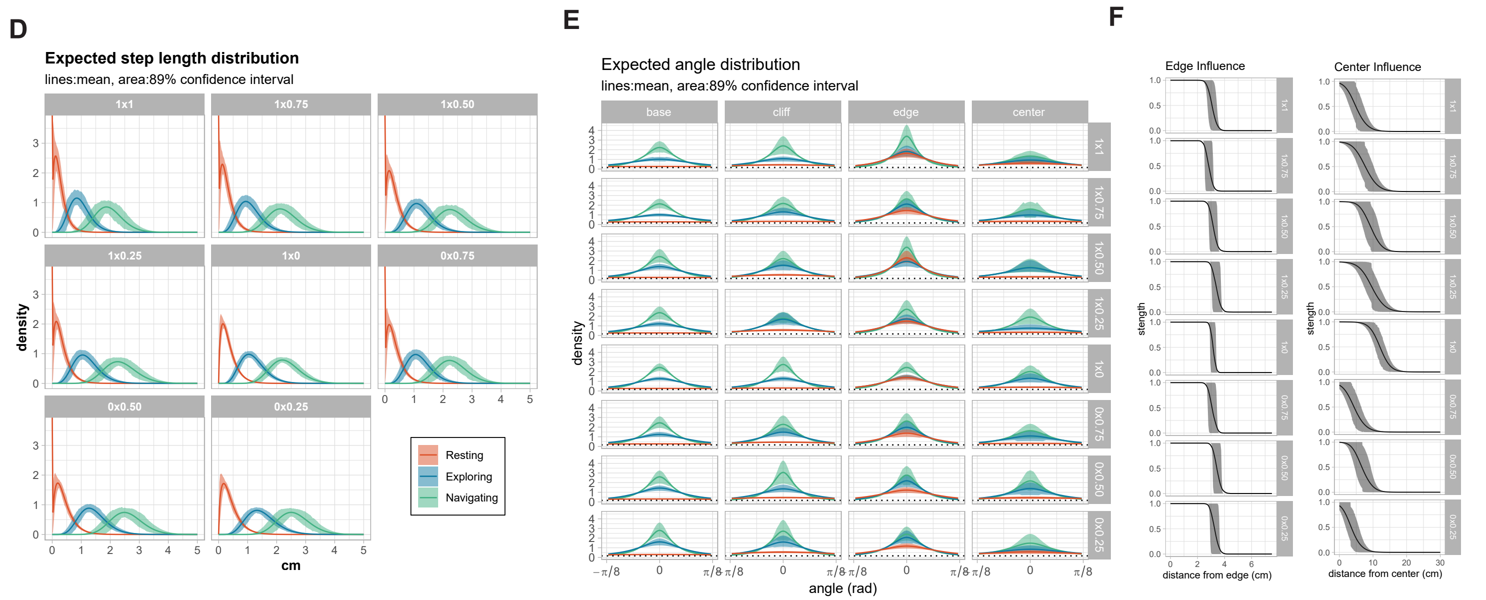


##### Time bin

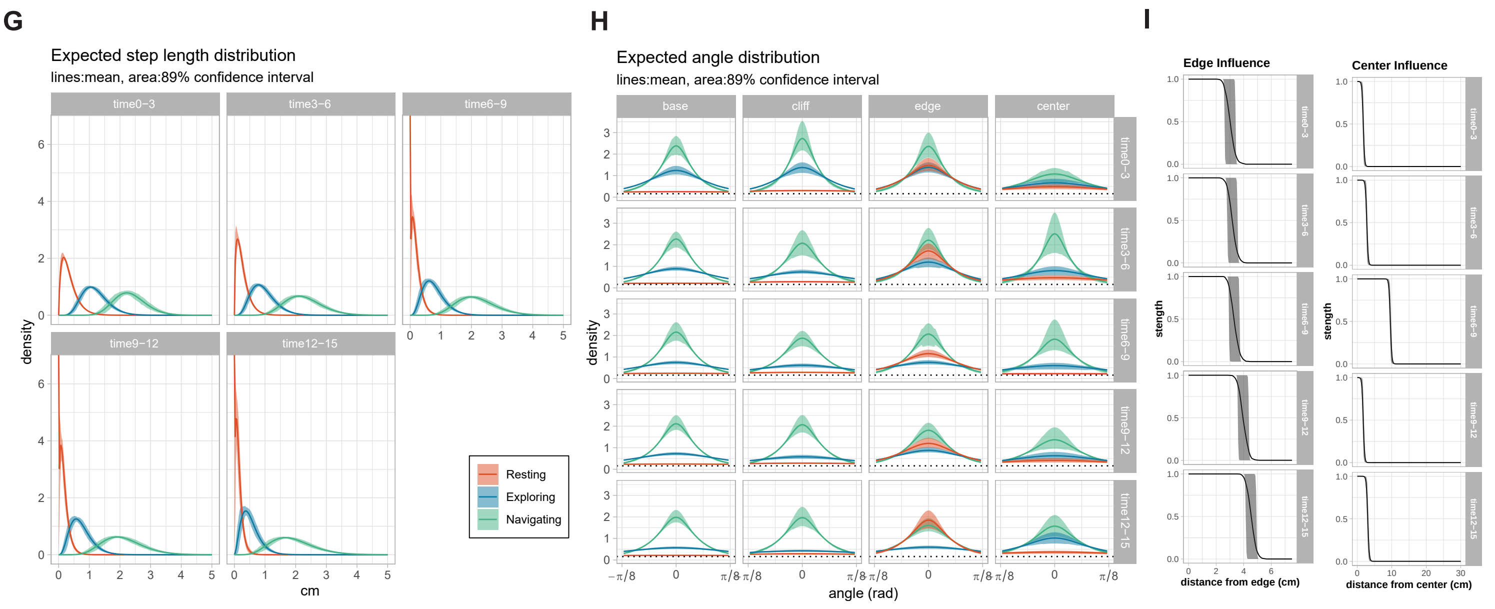


#### Supplemental Figure 5
